# Microbial Communities Performing Hydrogen Solventogenic Metabolism of Volatile Fatty Acids

**DOI:** 10.1101/2021.03.23.436570

**Authors:** Gustavo Mockaitis, Guillaume Bruant, Eugenio Foresti, Marcelo Zaiat, Serge R. Guiot

## Abstract

This work evaluates four different physicochemical pretreatments (acidic, thermal, acidic-thermal and thermal-acidic) on an anaerobic inoculum used for alcohol production from acetate and butyrate. All experiments were conducted in single batches using acetate and butyrate as substrates at 30 °C and with a pressurized headspace of pure H_2_ at 2.15 atm (218.2 MPa). Thermal and acidic-thermal pretreatments lead to higher production of both ethanol and butanol. Mathematical modelling shows that the highest attainable concentrations of ethanol and butanol produced were 122 mg L^-1^ and 97 mg L^-1^ for the thermal pretreatment (after 17.5 days) and 87 mg L^-1^ and 143 mg L^-1^ for the acidic-thermal pretreatment (after 18.9 days). Acetate was produced in all assays. Thermodynamic data indicated that a high H_2_ partial pressure favoured solventogenic metabolic pathways. Finally, sequencing data showed that both thermal and acidic-thermal pretreatments selected mainly the bacterial genera *Pseudomonas, Brevundimonas* and *Clostridium*. The acidic-thermal pretreatment selected a bacterial community more adapted to the conversion of acetate and butyrate into ethanol and butanol, respectively. Thermal-acidic pretreatment was unstable, showing significant variability between replicates. Acidic pretreatment showed the lowest alcohol production.

## 3 Background

The constant increase of prices of fossil fuels and the extensive land requirements for crop cultures targeting ethanol production is forcing the market to consider substitute avenues. The development of bioprocesses using organic residues as raw materials could be an alternative for fuel production. Anaerobic processes can be used to produce alcohol through solventogenic processes [1] and volatile fatty acids (VFAs) and hydrogen (H_2_) through acidogenic processes [2]. The conversion of any wastes into VFAs and H_2_ has an immediate commercial interest. Both propionic and butyric acids are raw materials of great interest, with many applications in various sectors, such as pharmaceutical and chemical industries [3]. H_2_ can be considered a raw material for subsequent processes and as an energy carrier to feed fuel cells [4].

Although anaerobic acidogenic processes could be used for wastes valorization, downstream processing of the VFAs obtained through those processes, such as bioconversion into their corresponding alcohols using H_2_ produced concomitantly, could have an even greater economic interest [5,6]. Ethanol and butanol produced through such processes can be used as drop-in liquid fuels, which have a higher market value per unit of energy. This two-steps approach (acidogenic followed by solventogenic processes) for ethanol and butanol production could both improve solvent production and reduce the toxicity linked to acidogenic processes products [6,7].

The most important parameters influencing anaerobic solventogenic fermentations include pH, organic acids, nutrient limitation, temperature, oxygen, and inoculum source [1,6,8]. Most of the studies performed on ethanol and butanol production through anaerobic solventogenic processes focussed on using sugars as carbon source and on using microbial pure cultures, with the most studied bacterial genera being *Thermoanaerobacter, Thermoanaerobacterium* and *Caloramator* under thermophilic conditions [9–13], and *Clostridium* [14–18].

Using mixed microbial cultures rather than pure cultures to produce alcohols from wastes through biological processes is of great interest. Mixed cultures increase process stability, improve the resistance to both toxicity and microbial contaminations, and bring higher substrate flexibility [19,20]. However, to date, there is still a lack of fundamental knowledge on mixed cultures used as inoculum, especially when H_2_ is used as an electron donor for the conversion of organic acids through solventogenic processes. Characterization of microbial communities capable of performing such processes is thus preponderant.

Enhancement of microbial communities through pretreatment of the inoculum is an effective way to induce changes in the communities to improve process performance. This approach has already been successfully tested, but mainly for optimizing the acidogenic step in H_2_ production through dark fermentation [21–23]. Pretreatments used in those studies consisted of modifications of the pH and temperature applied to the inoculum. Since microbial communities involved in acidogenic processes could use H_2_ as an electron donor to shift their metabolism to produce alcohols [1,5,24], the same inoculum pretreatments could be applied to improve solventogenic processes.

The present work evaluated the effects of four pretreatments of a mixed microbial population (acidic, thermal, acidic-thermal and thermal-acidic) on its capacity to convert VFAs into alcohols, using H_2_ as electron donor and an equimolar mixture of acetate and butyrate as carbon sources. The composition and dynamics of the mixed microbial community were analyzed.

## 4 Materials and Methods

### 4.1 Medium

The carbon source was an equimolar mixture of acetate and butyrate (17 mmol L^-1^, which correspond to 1,000 and 1,476 mg L^-1^ of acetate and butyrate, respectively). The nutrient medium (micro and macro) composition was prepared considering previous work on anaerobic digestion microbiology [25] and is described in Table 1. The initial pH was adjusted to 6.0.

**Table 1.** Nutrients composition of the medium [25].

| Nutrient | Concentration | Nutrient | Concentration | Nutrient | Concentration |
| --- | --- | --- | --- | --- | --- |
| $\text{NH}_4\text{Cl}$ | $1 \text{ g}\cdot\text{L}^{-1}$ | $\text{MnCl}_2\cdot 4\text{H}_2\text{O}$ | $50 \text{ }\mu\text{g}\cdot\text{L}^{-1}$ | $(\text{NH}_4)_6\text{Mo}_7\text{O}_{24}\cdot 4\text{H}_2\text{O}$ | $50 \text{ }\mu\text{g}\cdot\text{L}^{-1}$ |
| $\text{NaCl}$ | $100 \text{ mg}\cdot\text{L}^{-1}$ | $\text{AlCl}_3$ | $50 \text{ }\mu\text{g}\cdot\text{L}^{-1}$ | Pyridoxine Chloride | $10 \text{ }\mu\text{g}\cdot\text{L}^{-1}$ |
| $\text{MgCl}_2\cdot 6\text{H}_2\text{O}$ | $100 \text{ mg}\cdot\text{L}^{-1}$ | $\text{CoCl}_2\cdot 6\text{H}_2\text{O}$ | $50 \text{ }\mu\text{g}\cdot\text{L}^{-1}$ | HCl Concentrated | $1 \text{ }\mu\text{L}\cdot\text{L}^{-1}$ |
| $\text{CaCl}_2\cdot 2\text{H}_2\text{O}$ | $50 \text{ mg}\cdot\text{L}^{-1}$ | $\text{NiCl}_2\cdot 6\text{H}_2\text{O}$ | $92 \text{ }\mu\text{g}\cdot\text{L}^{-1}$ | $\text{Na}_2\text{SeO}_3\cdot 5\text{H}_2\text{O}$ | $100 \text{ }\mu\text{g}\cdot\text{L}^{-1}$ |
| K<sub>2</sub>HPO<sub>4</sub>·3H<sub>2</sub>O | 400 mg·L<sup>-1</sup> | EDTA | 500 µg·L<sup>-1</sup> | Nicotinic acid | 5 µg·L<sup>-1</sup> |
| FeCl<sub>2</sub>·4H<sub>2</sub>O | 2 mg·L<sup>-1</sup> | Biotin | 2 µg·L<sup>-1</sup> | Pantothenic acid | 5 µg·L<sup>-1</sup> |
| H<sub>3</sub>BO<sub>3</sub> | 50 µg·L<sup>-1</sup> | Riboflavin | 5 µg·L<sup>-1</sup> | B<sub>12</sub> Vitamin | 0.1 µg·L<sup>-1</sup> |
| ZnCl<sub>2</sub> | 50 µg·L<sup>-1</sup> | Thiamine | 5 µg·L<sup>-1</sup> | p-aminobenzoic acid | 5 µg·L<sup>-1</sup> |
| CuCl<sub>2</sub>·2H<sub>2</sub>O | 38 µg·L<sup>-1</sup> | Folic acid | 2 µg·L<sup>-1</sup> | Thioctic acid | 5 µg·L<sup>-1</sup> |

### 4.2 Raw Inoculum

The sludge used as raw inoculum in all assays was a primary digestate of the Carleton Corner Farms (Marionville, ON, Canada – 45º 11’ 14.0” N; 75° 21’ 54.1” W) collected in May 2013. The sludge was sieved three times using a 2 mm mesh sieve to eliminate all inert and heterogeneous lignocellulosic materials. The total volatile solids (TVS) content of the sieved sludge was 41 ± 3 mg TVS·L^-1^. The sludge was centrifuged (SorvalTM RC 6 Plus, Thermo Inc.) for 40 min at 10k min^-1^ and at 5°C. The supernatant was discarded, and the pellet was resuspended in a phosphate buffer (500 mg·L^-1^ PO_4_ ^3-^), using a homogenizer and disperser (Ultraturrax™ T25, IKA Inc.) for 10 min at 15k·min^-1^. The sludge was then sonicated to disaggregate possible granules and biofilms, using a sonicator (Vibra Cell™ VC130, Sonics Inc.) with 30W of power. This step was repeated four times, on ice, with a time/volume dependence relation of 4 s·mL^-1^, with 2 minutes interval between sonications. These steps were carried out to ensure the homogeneity of the inoculum and to wash possible organic dissolved materials present in the sludge, which could be used as an alternative carbon source during the process. The pH was corrected to 6.0 using a 1.0 M solution of HCl under vigorous stirring. This processed sludge was considered as the control inoculum.

### 4.3 Pretreatments

Four different physicochemical pretreatments – acidic, thermal, acidic-thermal, and thermal-acidic – were performed on the control inoculum. Pretreated inocula were then compared to each other, using the control inoculum as reference. All pretreated inocula were submitted to a starvation process to reduce the length of the lag phase before inoculation. This step consisted of incubating the inoculum for 72 hours at 30°C, under stirring of 50 min^-1^. The control inoculum was submitted to the same starvation process and presented a TVS content of 31 ± 0 mg TVS·L^-1^.

Acidic pretreatment consisted of decreasing the raw inoculum pH to 3.0 with a 12 M HCl solution under continuous stirring, followed by an incubation of 24 hours at 30°C, with stirring of 50 min^-1^. pH was then increased to 6.0 using a 2 M NaOH solution under stirring, followed by an incubation of 24 hours at 30°C under stirring of 50 min^-1^. In both pH decreasing and increasing inoculum, pH was controlled every hour for the five first hours of incubation to assure its stability. The acidic pretreated inoculum had a TVS of 29 ± 1 mg TVS·L^-1^.

The thermal pretreatment consisted of heating the control inoculum at 90°C for 20 minutes, under stirring and using a water batch. The inoculum was then immediately transferred to an ice batch until it reached room temperature (23°C). The thermally pretreated inoculum had a TVS of 31 ± 0 mg TVS·L^-1^.

The acidic-thermal and thermal-acidic pretreatments consisted of performing the two pretreatments sequentially, as indicated by the pretreatment name. The second pretreatment was performed immediately after the first one. The acidic-thermal pretreated inoculum had a TVS of 36 ± 3 mg TVS·L^-1^ and thermal-acidic pretreated inoculum had a TVS of 32 ± 0 mg TVS·L^-1^.

### 4.4 Experimental Setup

Initial physicochemical parameters for each experiment (pretreatments and control) and initial concentration of acetate, butyrate (before inoculation) and inoculum are shown in Table 2. Each experiment was carried out in quintuplicates, using 538 ± 3 mL sealed glass bottles, with an initial working volume of 110 mL. All the bottles were incubated upside down to avoid any diffusion of H_2_ through the bottle’s cap. All experiments were performed at 30°C under a 150 min^-1^ stirring. At the beginning of each experiment, the headspace of each bottle was totally replaced by pure H_2_ (99.99%) at a pressure of 2.39 ± 0.08 atm (242.2 ± 7.9 MPa). Such headspace replacement was repeated after each sampling to ensure constant pressure and composition all along with the experiment.

**Table 2.** Initial conditions (prior to inoculation) for each assay.

| Assay | pH | Acetate<br>mg·L<sup>-1</sup> | Butyrate<br>mg·L<sup>-1</sup> | Inoculum<br>mg TVS·L<sup>-1</sup> | Total H<sub>2</sub><br>mmol | Dissolved H<sub>2</sub><sup>*</sup><br>mmol·L<sup>-1</sup> |
| --- | --- | --- | --- | --- | --- | --- |
| Control | 5.89 | 1153 | 1727 | 11.3 ± 0.2 (5) | 37.9 ± 1.6 (5) | 1.69 |
| Acidic | 5.94 | 1139 | 1708 | 11.0 ± 0.2 (5) | 39.2 ± 0.2 (5) | 1.74 |
| Thermal | 6.07 | 1143 | 1712 | 12.8 ± 3.7 (5) | 40.2 ± 0.1 (5) | 1.78 |
| Acidic-thermal | 5.84 | 1233 | 1846 | 12.7 ± 0.2 (5) | 40.8 ± 0.2 (5) | 1.81 |
| Thermal-acidic | 5.85 | 1213 | 1816 | 11.3 ± 0.1 (5) | 39.7 ± 1.3 (5) | 1.77 |

### 4.5 Physicochemical Analyses

pH monitoring was performed with a portable potentiometer (Accumet™ AP115, Fisher Scientific™), with a microprobe electrode (Accumet™ 55500-40, Cole Parmer™), using the method 4500-H^+^ B, described by APHA [28]. TVS analyses were performed in quintuplicate, following method 2540-E, accordingly to APHA [28]. The pressure inside the bottles was measured using a digital manometer (DM8200, General Tools & Instruments™) with a range pressure of 0 - 6,804 atm (0 – 689.5 MPa). Dissolved CO_2_ was measured through alkalinity determination by potentiometric titration [29,30].

Mono, disaccharides, and organic fatty acids were analyzed using a Waters™ HPLC, which consisted of a pump (model 600) and an autosampler (model 717 Plus). The system was equipped with a refractive index detector (model 2414) for mono and disaccharides analyses. Organics acids were monitored from the same samples using the same equipment through a linked photodiode array detector (model 2996). A Transgenomic™ ICSep IC-ION-300 (300 mm x 7.8 mm outer diameter) column was used to separate all compounds and was operated at 35°C. The mobile phase was 0.01 N H_2_SO_4_ at 0.4 mL min^-1^ under an isocratic flow.

Alcohols (methanol, ethanol, acetone, 2-propanol, tert-butanol, n-propanol, sec-butanol, and n-butanol) were measured on an Agilent™ 6890 gas chromatograph (GC) equipped with a flame ionization detector [31].

100 µL gas samples, obtained using a gas-tight syringe (model 1750, Hamilton™), were used for gas composition (H_2_, CO_2_, CH_4_ and N_2_) measurements with a GC (HP 6890, Hewlett Packard™) equipped with a thermal conductivity detector (TCD) and an 11 m x 3.2 mm 60/80 mesh packed column (Chromosorb™ 102, Supelco™). The column temperature was held at 50 ºC for the entire run (4 min). The carrier gas was argon. The injector and detector were maintained at 125°C and 150°C, respectively.

### 4.6 16S rRNA gene Sequencing and microbial characterization

Bacterial 16S rRNA genes (V2 region) were amplified using the set of primers 16S-F343 IonA L01 (343-357; 5′ TACGGRAGGCAGCAG 3′) and 16S-R533 Ion P1 (516-533; 5′ ATTACCGCGGCTGCTGGC 3′) [32]. A sample-specific multiplex identifier was added to each forward primer and an Ion Torrent adapter to each primer. DNA extraction was conducted [33]. DNA was then purified [34,35]. Amplification reactions were performed in a final volume of 20 μL, which contained 1 μL of DNA, 0.5 μM of each primer, 7.5 μL of RNAse free H_2_O and 10 μL of 2X HotStarTaq™ Plus Master Mix (HotStarTaq™ *Plus* Master Mix Kit, Qiagen, USA). PCR conditions were an initial denaturation of 5 min at 95°C followed by 25 cycles of 30 s at 95°C, 30 s at 55°C, and 45 s at 72°C, with a final elongation step of 10 min at 72°C. PCR products were purified and quantified using a QIAquick™ gel extraction kit (Qiagen, USA) and a Quant-iT PicoGreen™ double-stranded DNA quantitation kit (Life Technologies Inc., USA) according to manufacturer’s instructions. The pooled amplicons were then sequenced using the Ion TorrentTM (Life Technologies Inc., USA) sequencing platform with a 314 chip. Bacterial 16S rRNA gene sequences generated were and analyzed using the ribosomal database project (RDP) classifier [36], using a bootstrap confidence cutoff of 50%, as recommended by RDP classifier [37] for short sequences (less than 250 bp). Before analysis, sequences shorter than 75 bp and sequences with unidentified bases (N) were removed.

### 4.7 Kinetics

Kinetic parameters for ethanol and butanol production were calculated using a modified Boltzmann sigmoidal model, as shown by Equation 1. The model was modified to incorporate parameter *r*_*max*_ as the maximum rate of the process (mg L^-1^ d^-1^).

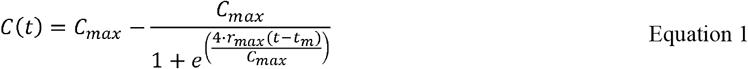

Where is *C*(*t*) the function of concentration in respect of time (mg L^-1^), *t* is time (d), *C*_*max*_ is the maximum concentration reached (mg L^-1^), and *t*_*m*_ is the value of time when 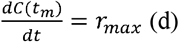.

From some of the calculated parameters showed in Equation 1, is possible to calculate the length of both lag (*t*_*i*_, d) and exponential (*t*_*e*_, d) phases, as depicted in Equation 3 and 4, respectively.

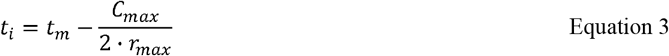

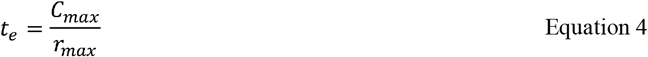

All fittings were performed using the software Microcal Origin Pro™ 9.0, using a Levenberg-Marquardt algorithm for fitting and initializing the equations parameters.

### 4.8 Metabolisms’ Gibb’s free energy

Gibb’s free energy values were calculated for all pathways depicted in Figure 1, except for pathway 1, for both initial (ΔG^I^_r_) and final (ΔG^F^_r_) conditions and were compared with the free energy for standard conditions (ΔG°_r_). Δ G°_r_ values for each reaction were estimated [38,39], considering 1 mol of each product and reagent at STP. ΔG^I^_r_ and ΔG^F^_r_ values were calculated using the Nernst equation at 30 °C and considered the concentrations of the metabolites presented in Table 4. To make all calculations possible, compounds that were not detected through the experiments, such as butanol, propionate, propanol, and ethanol, were assumed to be present, but at the same concentration of their respective lower detection limit for each methodology. This represents 10^−3^ mmol L^-1^ for volatile acids and alcohols and 10^−1^ mmol L^-1^ for CO_2_.

**Figure 1.**
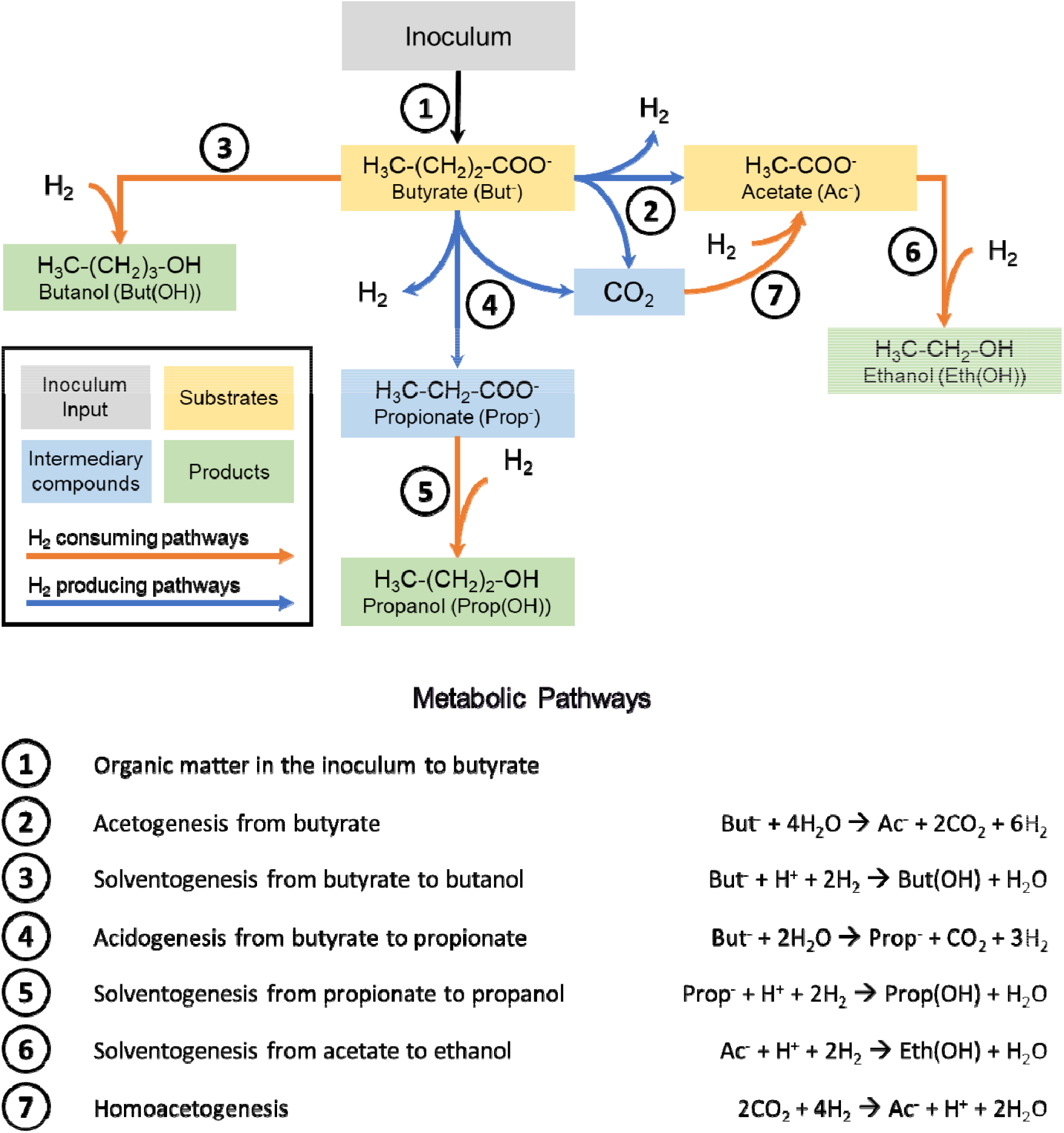
Alternative metabolic model proposed for the solventogenic process using butyrate and acetate as substrates. Pathways in orange are exergonic; pathways in blue are endergonic (at 30 °C, pH of 6.0, with a constant CO_2_ concentration of 10^−3^ mmol L^-1^, in initial conditions as depicted in Table 3).

## 5 Results and Discussion

### 5.1 Metabolic Model and Molar Balances

For all experiments, gas composition of the headspace was monitored prior to its replacement and H_2_ pressure was constantly kept at 2.15 ± 0.10 atm. H_2_ was the sole gas detected in the headspace in all cases, except for the control experiment, which showed a methane production of 0.24 mmol (5.37 mL at standard temperature and pressure (STP)). Initial and final concentrations of the main metabolites (added or produced) and of dissolved H_2_ and CO_2_ are presented in Table 3. Dissolved H_2_ was not measured but estimated through Henry’s law [26,27], and its concentration was considered constant, due to its continuous replacement. Dissolved CO_2_ was calculated from the alkalinity value measured at the beginning of each assay. CO_2_, O_2_ and N_2_ were never detected at the headspace. Acetone, lactate, and methanol were only sporadically detected in the liquid phase, in traces concentrations (below 1 mg L^-1^). Neither propionate nor alcohols were detected at the beginning of each experiment.

**Table 3.** Mean concentrations of all metabolites detected and of dissolved H_2_ and CO_2_.

| Metabolite<sup>a</sup> | Concentration (mg L<sup>-1</sup>) |  |  |  |  |  |  |  |  |  |
| --- | --- | --- | --- | --- | --- | --- | --- | --- | --- | --- |
|  | Control |  | Acidic |  | Thermal |  | Acidic-thermal |  | Thermal-acidic |  |
|  | Initial | Final | Initial | Final | Initial | Final | Initial | Final | Initial | Final |
| Acetate | 1225 ± 1 | 1575 ± 40 | 1222 ± 3 | 1492 ± 24 | 1206 ± 2 | 1722 ± 120 | 1235 ± 3 | 1645 ± 44 | 1224 ± 2 | 1447 ± 26 |
| Butyrate | 1831 ± 3 | 1773 ± 24 | 1828 ± 1 | 1935 ± 14 | 1791 ± 4 | 1796 ± 17 | 1840 ± 5 | 1664 ± 25 | 1831 ± 4 | 1738 ± 58 |
| Propionate | - | 42 ± 12 | - | 35 ± 4 | - | 107 ± 25 | - | 52.8 ± 9.2 | - | 40.1 ± 5.4 |
| Ethanol | - | 40 ± 6 | - | 14 ± 2 | - | 104 ± 7 | - | 94 ± 9 | - | 33 ± 6 |
| Propanol | - | - | - | 0.3 ± 0.2 | - | - | - | 1.8 ± 0.1 | - | 1.0 ± 0.3 |
| Butanol | - | 49 ± 8 | - | 11 ± 2 | - | 83 ± 3 | - | 154 ± 14 | - | 61 ± 25 |
| Dissolved H<sub>2</sub> | 1.6 ± 0.1 |  | 1.7 ± 0.0 |  | 1.6 ± 0.1 |  | 1.7 ± 0.1 |  | 1.7 ± 0.1 |  |
| Dissolved CO<sub>2</sub> | 3.07 | - | 2.40 | - | 3.30 | - | 1.56 | - | 2.19 | - |
| Duration (d) | 29.2 |  | 29.1 |  | 29.1 |  | 29.1 |  | 29.0 |  |

<sup>a</sup>All values are expressed in mg L<sup>-1</sup> except for dissolved H<sub>2</sub> and CO<sub>2</sub> expressed in mmol L<sup>-1</sup>. Mean concentrations of all volatile acids and alcohols was calculated considering 5 replicates. The amount of H<sub>2</sub> was kept constant in all experiments, there were no difference in its initial and final concentrations. Dissolved H<sub>2</sub> mean concentration considered 50 replicates.

Acetate and butyrate concentrations along the process demonstrated acetate production in all cases. Only minor variations of butyrate concentrations (slight decrease for the control and the acidic-thermal and thermal-acidic pretreatments, low production for the acidic and the thermal pretreatments) were observed. Such results suggested an alternative organic matter input in the system. This hypothesis is consistent with the decrease of the concentration of total volatile solids (TVS) that was observed for all conditions tested, as a direct effect of the different pretreatments on the biomass, which might have resulted in partial microbial cell death. Such non-living cells constituted non-soluble organic matter which was probably hydrolyzed and then consumed as a supplementary carbon source. Based on well-known anaerobic acidogenic and solventogenic metabolisms [1,24], and considering the inoculum as the only possible alternative source of organic matter, an alternative metabolic model including this alternative contribution and considering the metabolites (added and produced) presented in Table 3, was developed, as shown in Figure 1.

Figure 1 depicts butanol as the only product from lysed inoculum (pathway 1). Pathway 1 seems the only way to explain butanol production maintaining the butyric acid concentration constant since it is thermodynamically unfeasible to form butanol from acetate. Acetate could only be produced from butyrate, considering the high concentration of hydrogen and lack of CO_2_ in all assays, indicating that acetate was produced through acetogenesis from butyrate (pathway 2) or homoacetogenesis (pathway 7) after initial dissolved CO_2_ was consumed.

The metabolic model depicted in Figure 1 was used to perform a molar balance of all carbon inputs and outputs during the process (Table 4) and estimates the butyrate input from inoculum. All balances were performed on a 1 L basis and derived from initial and final experimental values obtained in all conditions (Table 3), except for the organic matter from the inoculum (pathway 1) which was expressed in butyrate equivalent.

**Table 4.** Molar balance of all metabolites detected in all metabolic pathways of each experiment.

| Metabolite | Pathway<sup>a</sup> | Molar Balance (mmol)<sup>b</sup> |  |  |  |  |
| --- | --- | --- | --- | --- | --- | --- |
|  |  | Control | Acidic | Thermal | Acidic-thermal | Thermal-acidic |
| Butyrate | Initial<sup>*</sup> | 21.0 | 21.0 | 20.6 | 21.1 | 21.0 |
|  | Final<sup>*</sup> | 20.4 | 22.2 | 20.6 | 19.1 | 20.0 |
|  | **1** | **+ 3.16** | **+ 3.56** | **+ 6.92** | **+ 4.73** | **+ 1.96** |
|  | 2 | - 2.53 | - 1.73 | - 4.34 | - 3.92 | - 1.57 |
|  | 3 | - 0.66 | - 0.15 | - 1.12 | - 2.08 | - 0.82 |
|  | 4 | - 0.57 | - 0.48 | - 1.46 | - 0.75 | - 0.57 |
| Butanol | 3<sup>*</sup> | + 0.66 | + 0.15 | + 1.12 | + 2.08 | + 0.82 |
| Propionate | 4<sup>*</sup> | + 0.57 | + 0.48 | + 1.46 | + 0.75 | + 0.57 |
|  | 5 | - | - | - | - 0.03 | - 0.02 |
| Propanol | 5<sup>*</sup> | - | - | - | + 0.03 | + 0.02 |
| Acetate | Initial<sup>*</sup> | 20.7 | 20.7 | 20.4 | 20.9 | 20.7 |
|  | Final<sup>*</sup> | 26.7 | 25.3 | 29.2 | 27.9 | 24.5 |
|  | 2 | + 2.53 | + 1.73 | + 4.34 | + 3.92 | + 1.57 |
|  | 6 | - 0.87 | - 0.30 | - 2.26 | - 2.04 | - 0.72 |
|  | 7 | + 4.35 | + 3.17 | + 6.72 | + 5.08 | + 2.95 |
| Ethanol | 6<sup>*</sup> | + 0.87 | + 0.30 | + 2.26 | + 2.04 | + 0.72 |
| Dissolved CO<sub>2</sub> | Initial<sup>*</sup> | 3.07 | 2.40 | 3.30 | 1.56 | 2.19 |
|  | 2 | + 5.05 | + 3.46 | + 8.68 | + 7.85 | + 3.14 |
|  | 4 | + 0.57 | + 0.48 | + 1.46 | + 0.75 | + 0.57 |
|  | 7 | - 8.69 | - 6.34 | - 13.4 | - 10.2 | - 5.90 |

<sup>\*</sup>Values obtained experimentally (derived from initial and final concentrations as indicated in Table 3).
<sup>a</sup>Metabolic pathways described in Figure 1.
<sup>b</sup>Balance on a 1 L basis; “+” indicates a production; “-” indicates a consumption.

According to the mass balance shown in Table 4, CO_2_ was produced by converting butyrate into acetate (pathway 2) and by acidogenesis from butyrate to propionate (pathway 4). This CO_2_ was then consumed to form acetate through a homoacetogenic pathway (pathway 7). This hypothesis was supported by the absence of gaseous CO_2_ in all experiments, followed by an increase in acetate concentration. This CO_2_ absence may be linked to microbial communities more adapted to convert butyrate and CO_2_ into acetate (pathways 2 and 7, respectively) rather than to the conversion of acetate into ethanol (pathway 6). This metabolic model also indicates that butyrate was mainly consumed to produce acetate (pathway 2) in all studied conditions and the organic matter from the inoculum (pathway 1) represented an external input since butyrate concentrations were almost constant in all assays, although consumed to form butanol (pathway 3). Ethanol and butanol were produced (pathways 3 and 6) in all experiments. These pathways were more active in the experiments with thermal and acidic-thermal pretreatments, as these conditions showed the highest ethanol and butanol production among all other experiments. Although acidogenesis from butyrate to propionate (pathway 4) was active in all conditions, conversion of propionate into propanol (pathway 5) was not an important metabolic pathway. Propanol production was not expressive in neither experiments since it showed the highest concentration of 1.82 mg L^-1^ in acidic-thermal pretreatment essay.

The successful closure of molar balances shown in Table 4, with a stoichiometrically balanced sum of all inputs and outputs (considering the contribution of the inoculum), indicates that the metabolic model proposed in Figure 1 accurately represents the solventogenic processes occurring in all studied conditions. An energy balance based on the mass balance depicted in Table 4 was calculated to estimate Gibb’s free energy values for each pathway for both initial and final conditions (Figure 2). Those values provide an overview of the thermodynamic feasibility of each pathway involved in the solventogenic process and tend to validate the metabolism proposed in Figure 1.

**Figure 2.**
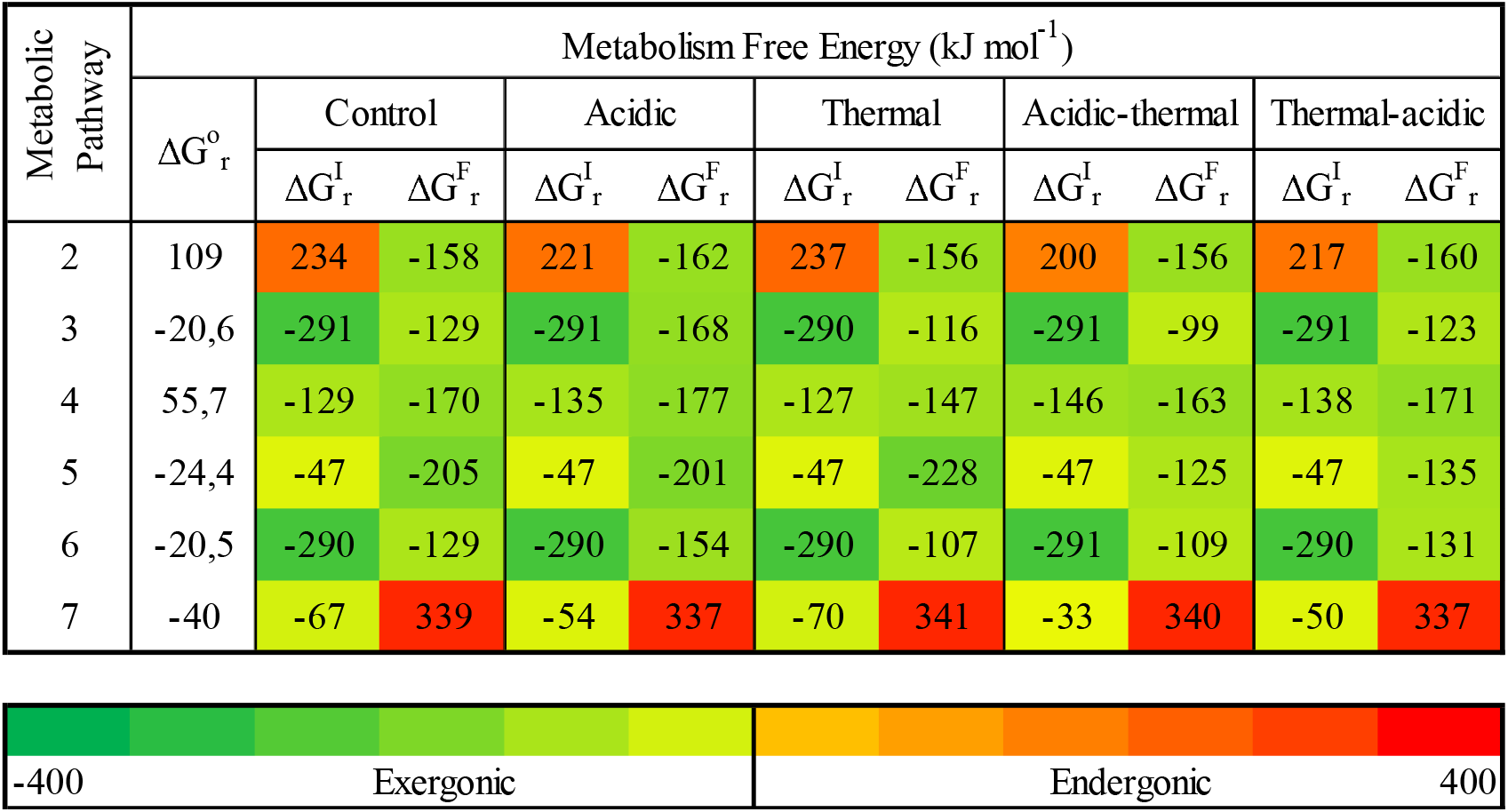
Gibbs’ free energy for initial (ΔG^I^ _r_), final (ΔG^F^_r_) and standard (ΔG°_r_) conditions for each pathway (as defined in Figure 1) in each experiment. Greenish values (< 0) are exergonic and reddish values (> 0) are endergonic.

Energetic profiles could help to anticipate changes within metabolic pathways. As shown in Figure 2, the estimated ΔG°_r_ values indicate that acetogenesis from butyrate and acidogenesis from butyrate to propionate (pathways 2 and 4, respectively) are theoretically thermodynamically unfeasible metabolisms. Nevertheless, these pathways became thermodynamically feasible due to the absence (or low concentration) of dissolved CO_2_ at the course of each essay. Acetogenesis from butyrate (pathway 2) was likely inhibited at the beginning of each experiment due to the concentration of H_2_ and CO_2_. During the experiment, CO_2_ was progressively consumed through homoacetogenesis (pathway 7), which rendered pathway 2 thermodynamically feasible. At low concentrations of CO_2_ as found after the beginning of each essay, the homoacetogenic pathway might have become thermodynamically unfeasible. Gibb’s free energy values found for these pathways could explain the production of acetate observed in all studied conditions, all along the process. Metabolic pathways 3 (solventogenesis from butyrate to butanol) and 6 (solventogenesis from acetate to ethanol) were thermodynamically feasible in all studied conditions all along the process, explaining the observed production of such alcohols. Metabolic pathway 4 (acidogenesis from butyrate to propionate) was unfeasible at the ending of each condition, showing that propionate production was stationed in a low concentration at some point of the experiments.

As described previously, H_2_ has a significant role as a co-substrate in the anaerobic process energetics [40,41]. As shown in Figure 3, solventogenesis of butanol from butyrate (pathway 3) is less sensitive to low ppH_2_ than solventogenesis of ethanol and propanol, respectively from acetate (pathway 6) and propionate (pathway 5). By extrapolation, equilibrium (ΔG°_r_ = 0) in pathway 3 would be achieved at a ppH_2_ of 4·10^−4^ atm (43.7 Pa), implying that solventogenesis of butanol could theoretically be carried out at very low ppH_2_. It is also possible to infer from Figure 3 that at ppH_2_ higher than 0.62 atm (62.5 MPa), acidogenesis of propionate (pathway 4) and acetogenesis of acetate (pathway 2) from butyrate will stop, shutting down the production of acetate, propionate and H_2_. In addition to low CO_2_ concentrations, high ppH_2_ thus represents an important factor that renders such process thermodynamically feasible and favours solventogenesis from VFAs.

**Figure 3.**
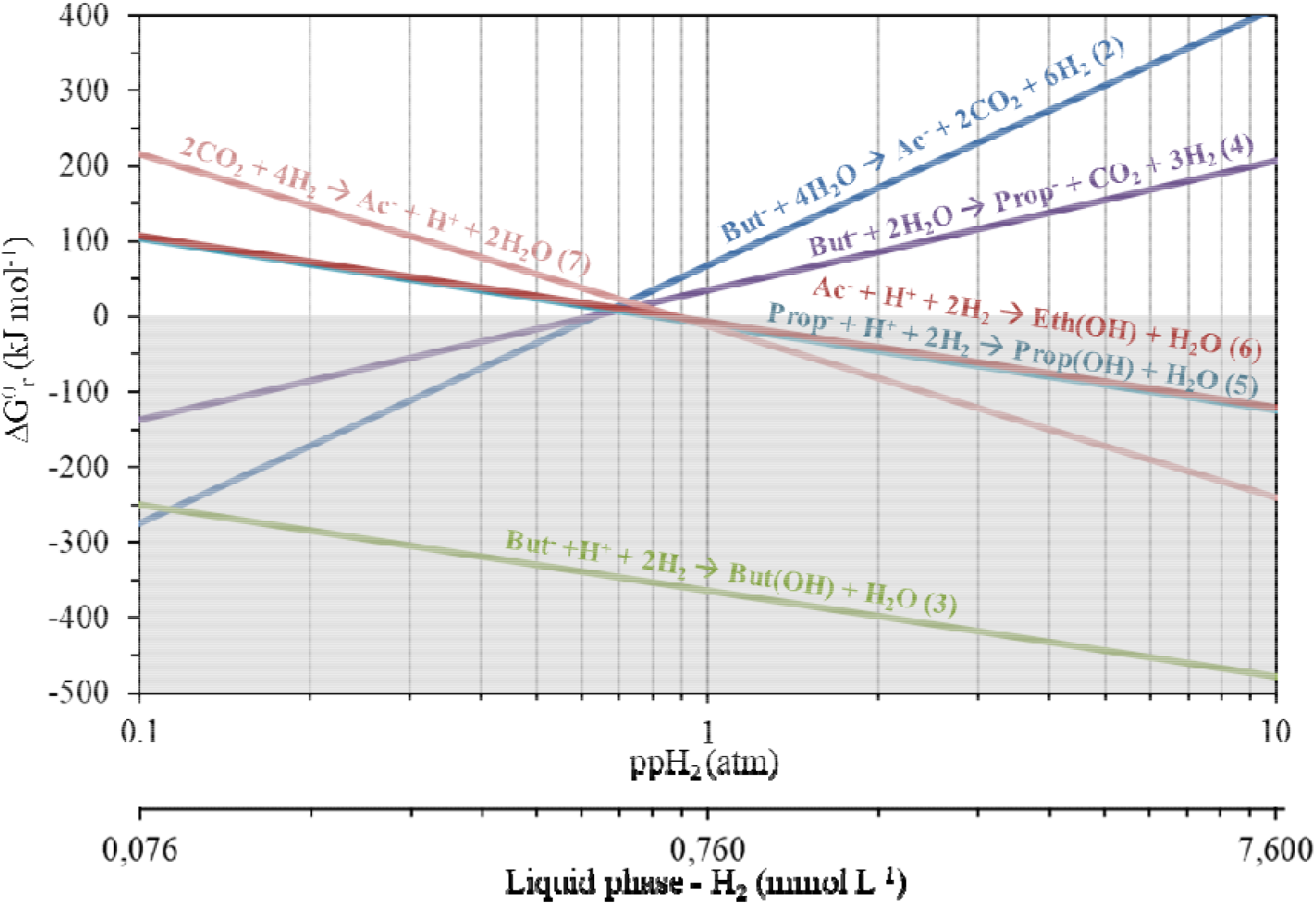
Evolution of the standard Gibbs’ free energy (ΔG°_r_) in function of the partial pressure of H_2_. Evolution for each metabolic pathway (as defined in Figure 1), at 30 °C, and with a standard concentration of 1.0 mol L^-1^ for each reagent other than H_2_. Shadowed area indicates thermodynamically feasible reactions.

### 5.2 Alcohols and VFAs metabolisms

As shown in Figure 4, the highest concentrations of alcohols were obtained for both thermal and acidic-thermal pretreatments, with the best rate observed for the acidic-thermal pretreatment, especially for butanol production. On the opposite, the lowest alcohol production was observed for the acidic pretreatment. Acetate and butyrate concentrations along the essays confirmed the contribution of the inoculum as an important source of organic matter as the substrate for alcohol production, since in all conditions tested, acetate concentration increased, and butyrate concentration increased or only slightly decreased. These results reinforced the hypothesis described in Figure 1 and Table 4 of an acetate production through acetogenesis of butyrate (pathway 2) and homoacetogenesis (pathway 7), and preferentially through pathway 2 due to low concentrations of CO_2_ at the beginning of all assays.

**Figure 4.**
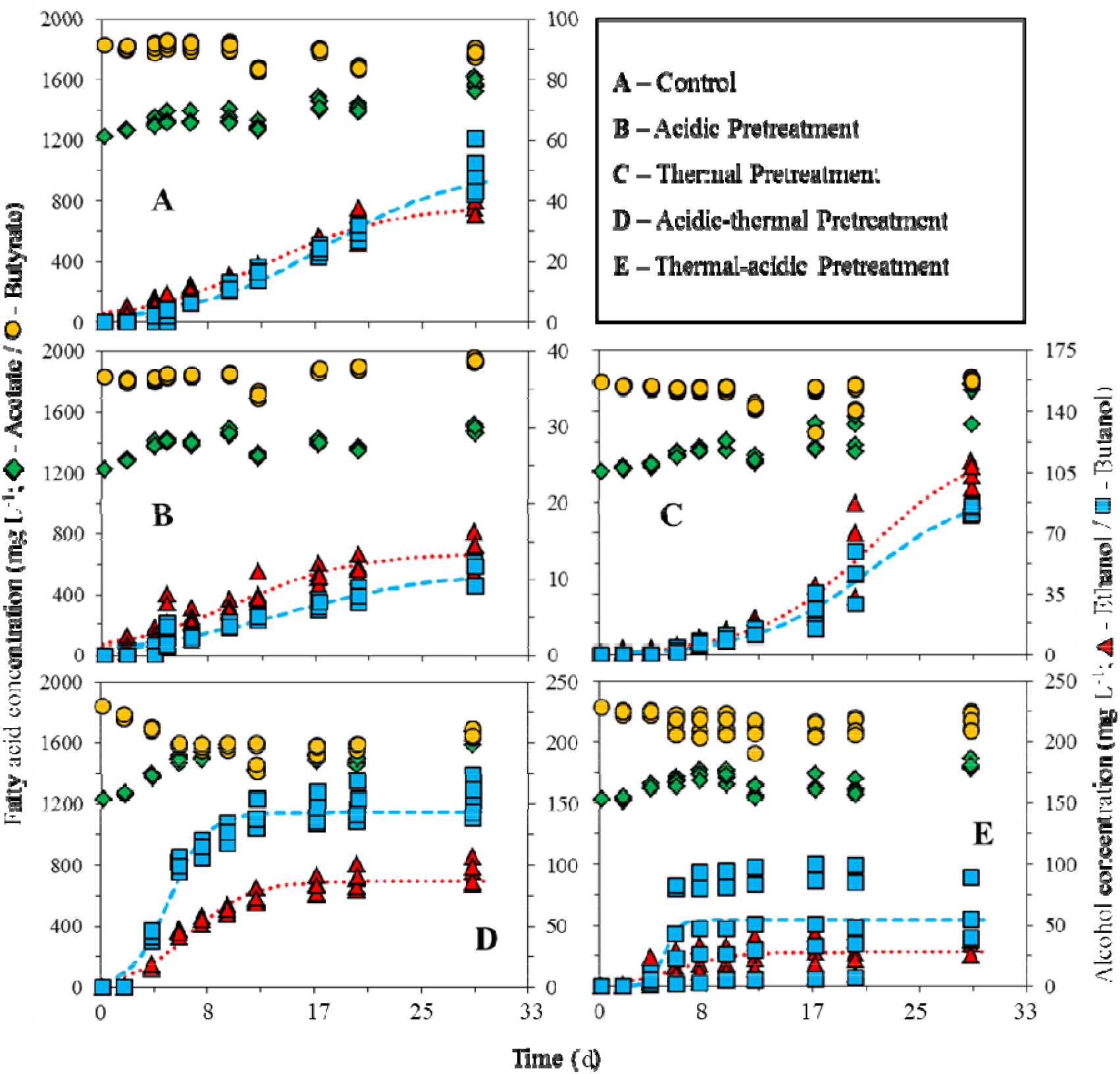
Kinetics of VFAs (acetate and butyrate) and alcohols (ethanol and butanol) productions for each studied condition. Red dotted- and blue dashed-lines represent the modified Boltzmann model fittings for ethanol and butanol concentrations, respectively.

As shown in Figure 5, propionate was produced in all conditions tested. No significant difference was observed in the concentrations produced (42.4 ± 10.0 mg L^-1^), except for the thermal pretreatment essay, for which the level of production was 2.5 times higher than in all other conditions. As preseted in Table 3, propionate was then consumed and converted into propanol (pathway 5, Figure 1) for the acidic, acidic-thermal and thermal-acidic pretreatments. Despite the favourable thermodynamics of this reaction (Figure 2), propanol was only produced in trace amounts. Oxidation of propionate into acetate was considered unlikely since such reaction is energetically unfeasible in standard conditions (ΔG°_r_ = 53.3 kJ mol^-1^), and that, as previously described [39], at high H_2_ concentrations ΔG_r_ values are increased.

**Figure 5.**
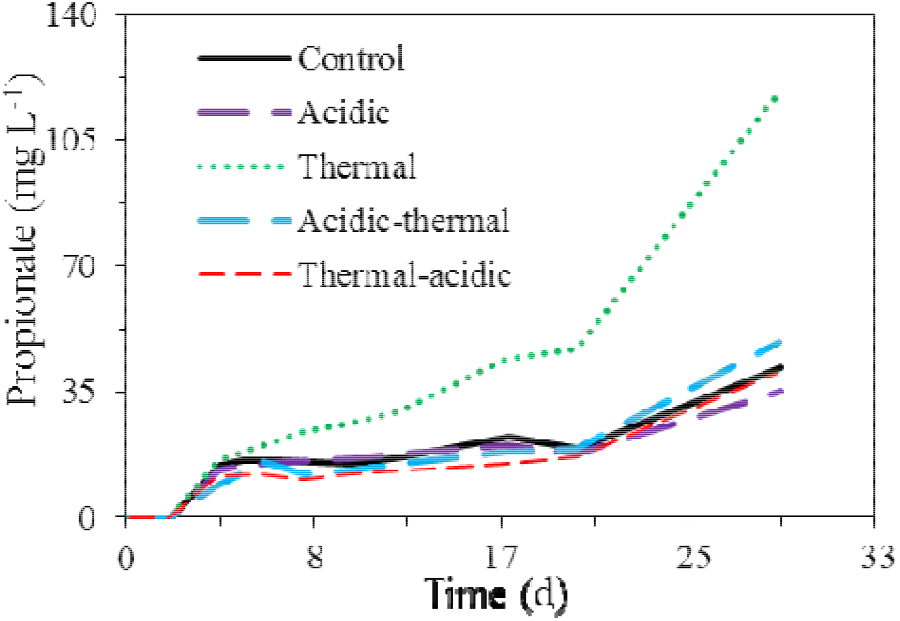
Behaviour of propionate concentration for each studied condition.

Table 5 presents the parameters of the modified Boltzmann model fitting the experimental data from Figure 4. Those parameters are related to the kinetics of alcohol production and compare efficiencies between all pretreatments. A high correspondence was obtained between replicates for all conditions tested, apart from the thermal-acidic pretreatment. The correlation coefficient was very low in that condition, only reaching 0.5 for ethanol and 0.4 for butanol, indicating that the process was unpredictable and could not be reproduced. Due to its instability and unpredictability, the thermal-acidic pretreatment was thus no longer considered for analyses.

**Table 5.**
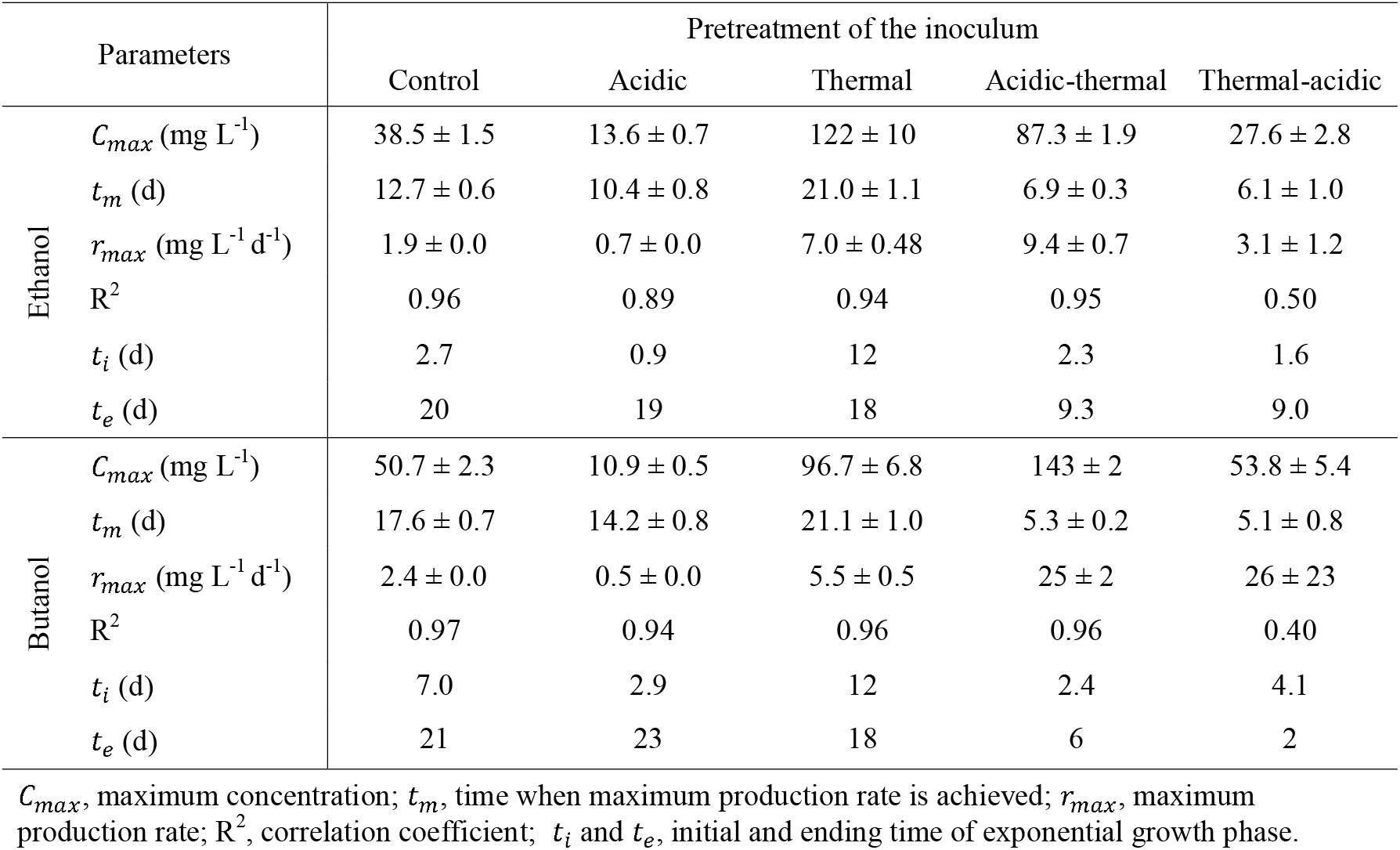
Parameters of the modified Boltzmann model fitting for ethanol and butanol production.

| Parameters |  | Pretreatment of the inoculum |  |  |  |  |
| --- | --- | --- | --- | --- | --- | --- |
|  |  | Control | Acidic | Thermal | Acidic-thermal | Thermal-acidic |
| Ethanol | $C_{max}$ (mg L<sup>-1</sup>) | 38.5 ± 1.5 | 13.6 ± 0.7 | 122 ± 10 | 87.3 ± 1.9 | 27.6 ± 2.8 |
| | $t_m$ (d) | 12.7 ± 0.6 | 10.4 ± 0.8 | 21.0 ± 1.1 | 6.9 ± 0.3 | 6.1 ± 1.0 |
| | $r_{max}$ (mg L<sup>-1</sup> d<sup>-1</sup>) | 1.9 ± 0.0 | 0.7 ± 0.0 | 7.0 ± 0.48 | 9.4 ± 0.7 | 3.1 ± 1.2 |
|  | R<sup>2</sup> | 0.96 | 0.89 | 0.94 | 0.95 | 0.50 |
| | $t_i$ (d) | 2.7 | 0.9 | 12 | 2.3 | 1.6 |
| | $t_e$ (d) | 20 | 19 | 18 | 9.3 | 9.0 |
| Butanol | $C_{max}$ (mg L<sup>-1</sup>) | 50.7 ± 2.3 | 10.9 ± 0.5 | 96.7 ± 6.8 | 143 ± 2 | 53.8 ± 5.4 |
| | $t_m$ (d) | 17.6 ± 0.7 | 14.2 ± 0.8 | 21.1 ± 1.0 | 5.3 ± 0.2 | 5.1 ± 0.8 |
| | $r_{max}$ (mg L<sup>-1</sup> d<sup>-1</sup>) | 2.4 ± 0.0 | 0.5 ± 0.0 | 5.5 ± 0.5 | 25 ± 2 | 26 ± 23 |
|  | R<sup>2</sup> | 0.97 | 0.94 | 0.96 | 0.96 | 0.40 |
| | $t_i$ (d) | 7.0 | 2.9 | 12 | 2.4 | 4.1 |
| | $t_e$ (d) | 21 | 23 | 18 | 6 | 2 |

$C_{max}$ , maximum concentration; $t_m$ , time when maximum production rate is achieved; $r_{max}$ , maximum production rate; R<sup>2</sup>, correlation coefficient; $t_i$ and $t_e$ , initial and ending time of exponential growth phase.

As shown in Figure 4 and Table 5, both alcohol production and maximum production rates (*r*_*max*_) were improved for thermal and acidic-thermal pretreatments. Although the highest ethanol production was observed for the thermal pretreatment, the best results were obtained for the acidic-thermal pretreatment, which allowed the best butanol production and an increase of 4.5 times of the ethanol and of 10.2 times of the butanol maximum production rate. In opposition, a decrease in alcohol production was observed for the acidic pretreatment. Such results clearly indicate that an acidic-thermal pretreatment of the inoculum is a promising approach for designing more efficient and smaller-sized bioreactors.

Length of the lag phase (*t*_*i*_) and duration of the bacterial exponential growth phase (*t*_*e*_) were evaluated for each pretreatment and compared to the control by considering alcohol production curves as growth-associated curves (Table 3). The length of the lag phase is related to the time required for a bioreactor to initiate its process (start-up) and achieve higher rates of alcohol production. For both ethanol and butanol production, the shortest lag phases were observed for the acidic and acidic-thermal pretreatments. In both pretreatments, the lag phase was shorter than found in the control essay. The most extended lag phase was observed for the thermal pretreatment, which was considerably greater for both ethanol and butanol production than observed in control. Such results indicate that the thermal pretreatment had the highest impact on the inoculum, while acidic and acidic-thermal pretreatments selected microbial communities which were the best adapted to solventogenic processes. Variations in the duration of the bacterial exponential growth phases (*t*_*e*_) were also observed (Table 5). A shorter exponential growth phase is representative of a faster process. However, this parameter must be evaluated concomitantly with the maximum rates of production (*r*_*max*_). For example, a short *t*_*e*_ occurring at low *r*_*max*_ indicates a process occurring with low efficiency. Among all the conditions tested, low *t*_*e*_ values and high *r*_*max*_ values were obtained for the acidic-thermal pretreatment for ethanol and butanol production. Taken all together, results obtained for all conditions tested indicate that the acidic-thermal pretreatment was the best approach to generate a rapid and efficient process.

pH was initially set to a value of 5.92 ± 0.09 for all essays, and their value increased during the experiments (Figure 6). This rise in pH values likely reflects the consumption of H2 as an electron donor to form alcohols since solventogenic pathways (3, 5 and 6) require the consumption of H+ and alcohols show a low ionization on water. Such a pH increase occurred to a lesser extent in all conditions, including an inoculum acidic pretreatment (acidic, acidic-thermal and thermal-acidic pretreatments). This difference might be explained by a more drastic initial pH drop resulting from the acidic addition on these pretreatments, probably lowering the buffering capacity of the inoculum. This hypothesis is reinforced by the concentrations of dissolved CO_2_, which were higher for both control and thermal pretreatment than acidic and acidic-thermal pretreatments (Table 3).

**Figure 6.**
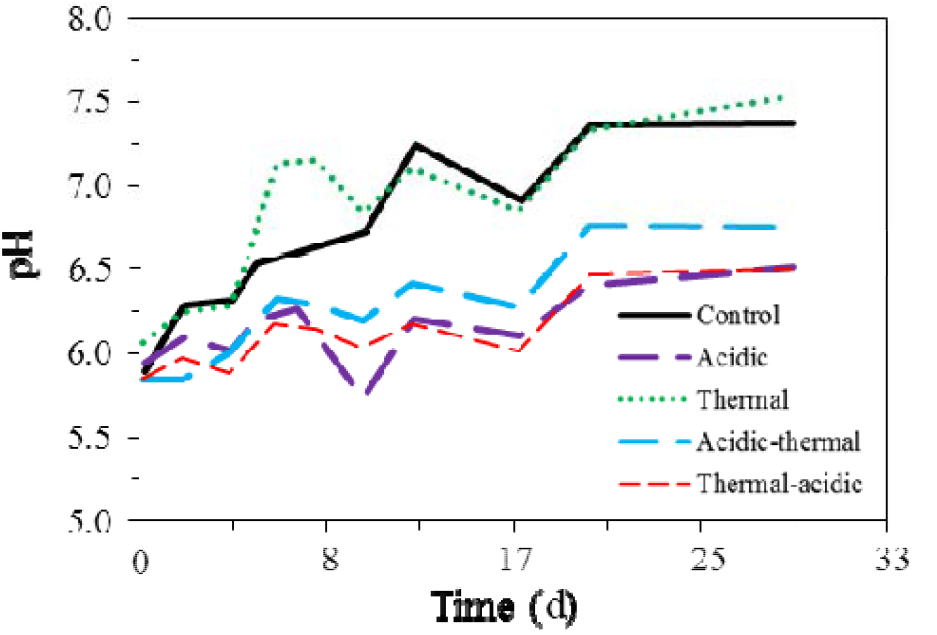
Evolution of pH along the process for each studied condition.

A decrease in total volatile solids (TVS) was also observed for all conditions tested (Figure 7). This decrease occurred probably due to a direct effect of the different pretreatments on the biomass, resulting in partial microbial cell death. Such non-living cells constituted non-soluble organic matter which was probably hydrolyzed and then consumed as a carbon source, contributing to the organic matter input proposed in pathway 1 of Figure 1. The TVS concentration stopped decreasing and started to increase after 6.3 days in the acidic-thermal pretreatment slightly. Such evolution is probably linked to the higher values of observed for this pretreatment (Table 5), which reflect an increase in the biomass growth’s rate, and consequently, the TVS concentration.

**Figure 7.**
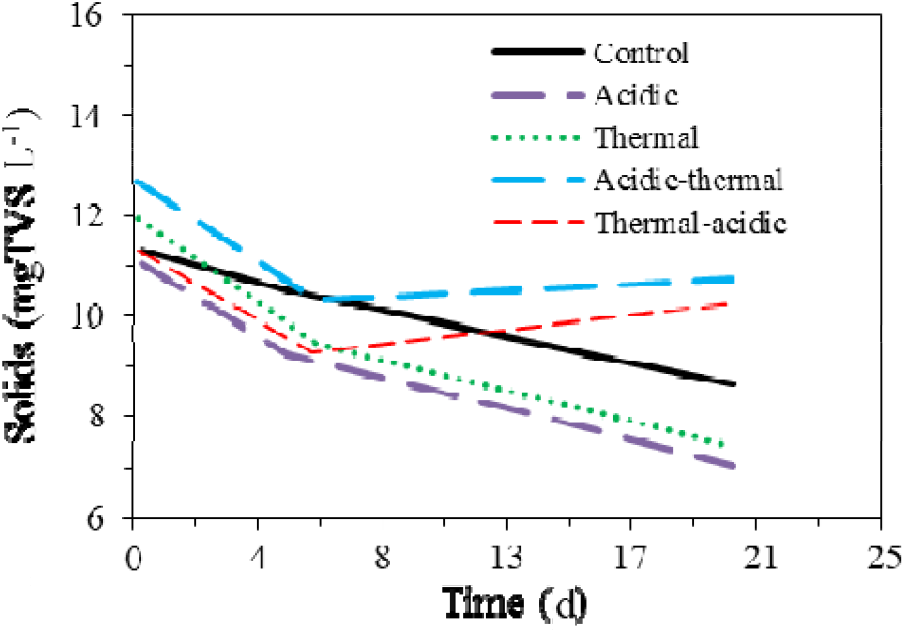
Evolution of the total volatile solids (TVS) along the process for each studied condition.

### 5.3 Bacterial communities

Based on the results presented above and focusing on alcohol production, both thermal and acidic-thermal pretreatments experiments were submitted to microbial community analyses. Bacterial populations were thus characterized and monitored all along with those two processes. Thirteen phyla were detected, namely *Firmicutes, Proteobacteria, Bacteroidetes, Actinobacteria, Synergistetes, Cyanobacteria, Tenericutes, Spirochaetes, Deinococcus-Thermus, Fibrobacteres, Verrucomicrobia, Chloroflexi* and *Nitrospirae*. Among those, three (*Firmicutes, Proteobacteria* and *Bacteroidetes*) represented up to 94.6% of the total OTUs detected in each sample for both pretreatments (Figure 8). *Firmicutes* was the most abundant phylum detected in the initial non-treated inoculum (corresponding to 66.6% of the detected OTUs). Both pretreatments generated a shift in the bacterial population by stimulating the development of *Proteobacteria* and strongly reducing the number of *Firmicutes*. Bacterial populations evolved differently along the process for the two pretreatments. In the thermal pretreatment experiment (Figure 8A), a decrease of *Proteobacteria* was observed concomitantly with an increase in *Bacteroidetes* and *Firmicutes*, the latest becoming the dominant phylum at the end of the process. This shift started after 200 hours of experiment, which correspond to the early beginning of the bacterial exponential growth phase (Figure 4C and Table 5). In opposition, in the acidic-thermal pretreatment experiment (Figure 8B), relative stability was observed for the bacterial population, with *Proteobacteria* remaining the main phylum all along the process (72.0 ± 4.99% of the detected OTUs) and *Firmicutes* constantly being the second phylum of importance (24.0 ± 5.10% of the detected OTUs). *Bacteroidetes* increased during the process, before starting to decrease after 500h of experiment, to reach their initial level at the end of the experiment.

**Figure 8.**
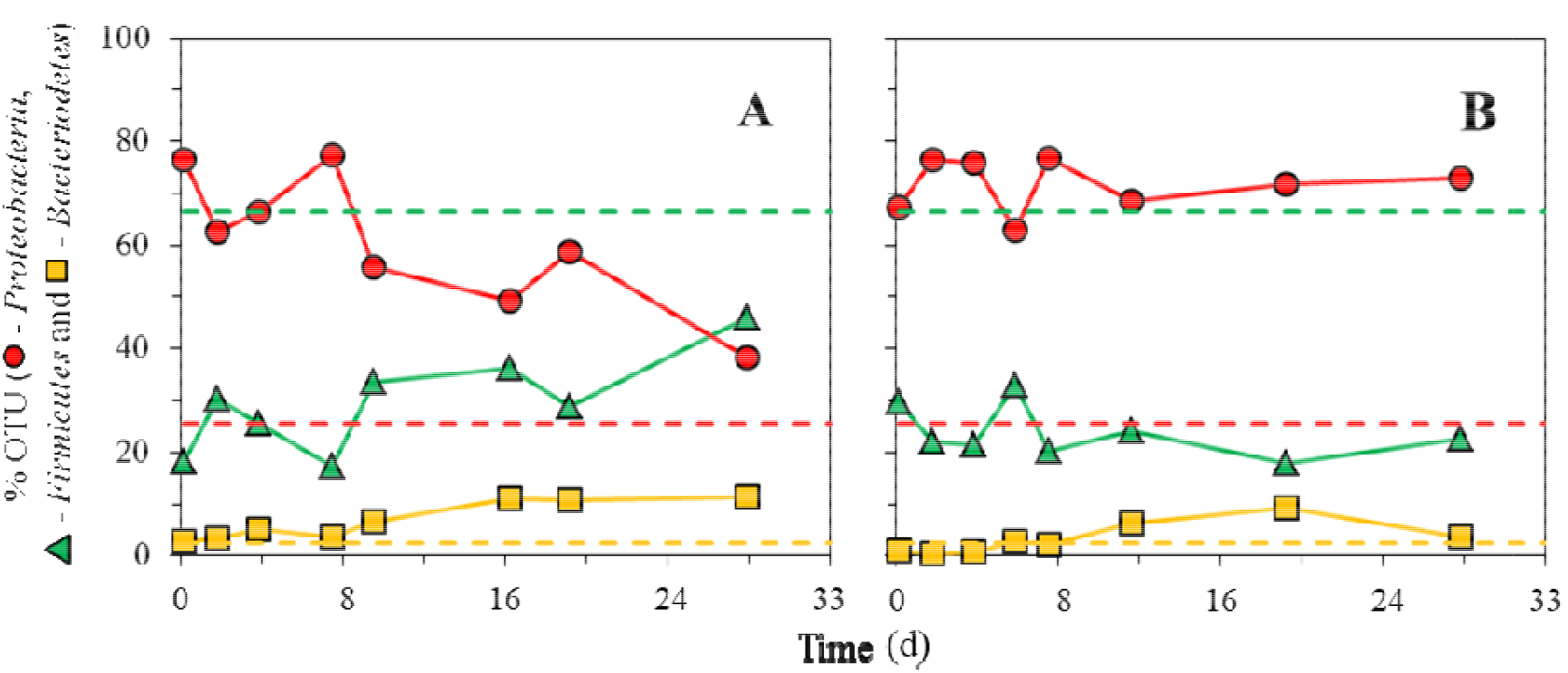
Evolution of the three most representative bacterial phyla for the thermal (A) and acidic-thermal (B) pretreatments. Dashed lines represent OTUs associated to those same phyla found in the inoculum without pretreatment.

Deeper phylogenetic analyses were performed down to the genus level to infer the potential metabolic pathway(s) that could be associated with the enhancement of alcohol production observed for both thermal and acidic-thermal pretreatments (Figure 9). Substantial differences were noticed between the bacterial population from the initial inoculum (before any pretreatment) and those from the thermal and acidic-thermal pretreated inocula.

**Figure 9.**
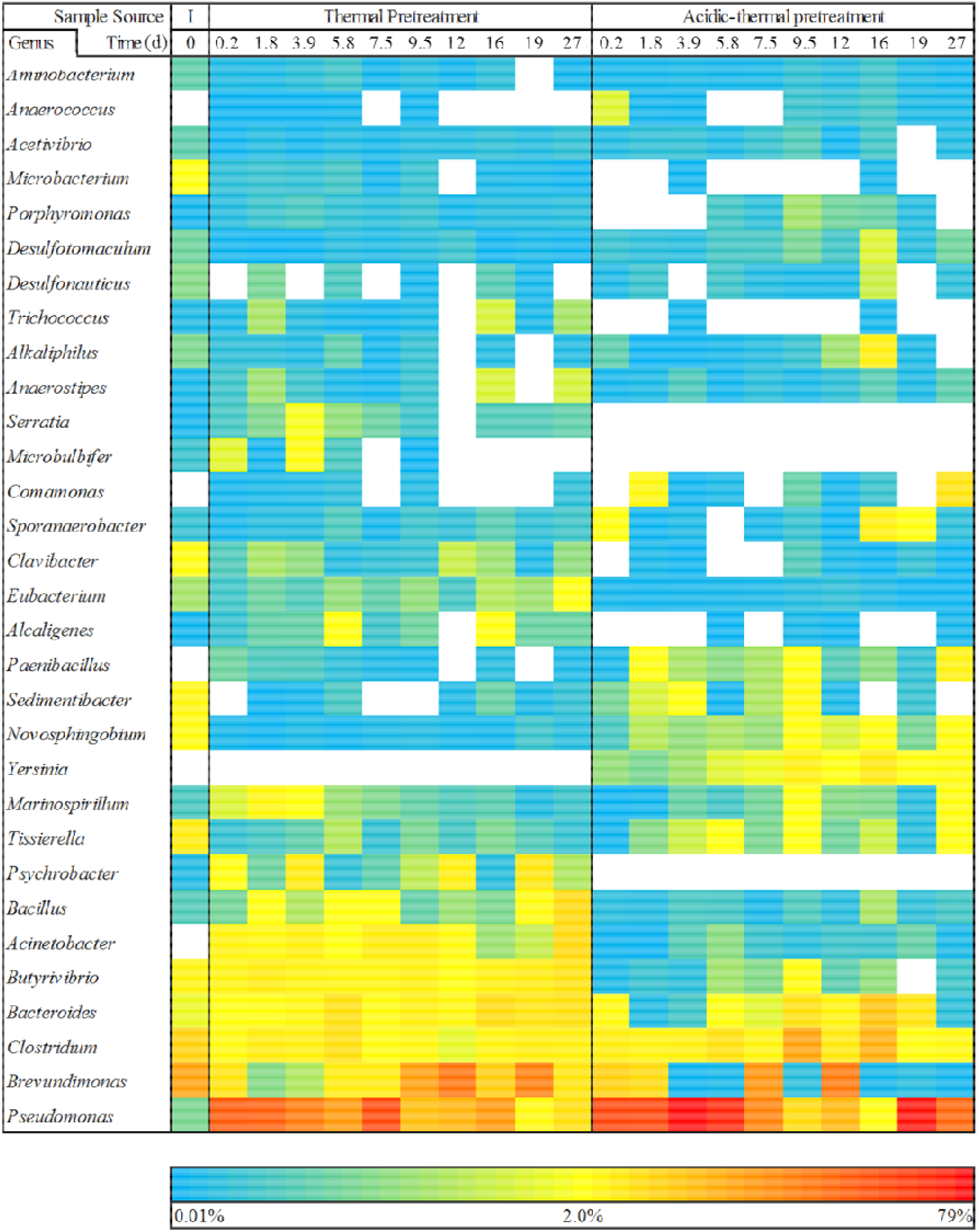
Evolution of the bacterial population at the genus taxonomic level for thermal and acidic-thermal pretreatments, and comparison with the initial bacterial population (I) for the 31 most detected genera.

A total of 317 genera were detected in all samples tested, but only 31 of them, representing up to 85% of the bacterial population, were considered for further analysis (Figure 9). One of the most noticeable changes consisted of an increase of *Pseudomonas* in pretreatment experiments compared to the initial population. Such increase was observed as soon as the first hours of experiments. *Pseudomonas* then remained a dominant genus of the bacterial population, despite a slight decrease observed for the thermal pretreatment in the second half of the process. Several *Pseudomonas* species have been genetically well-characterized and have been shown to possess the genetic components for both ethanol (from pyruvate) and butanol (from glycerol and pyruvate) production [42]. The prevalence of *Pseudomonas* is likely linked with the higher levels of alcohol production observed in thermal and acidic-thermal pretreatments. Some *Pseudomonas* species also possess genes responsible for the degradation of ethanol into acetyl-CoA [42]. A pathway related to ethanol degradation might explain better butanol production compared to ethanol, observed for the acidic-thermal pretreatment. Among the other noticeable results, two bacterial genera, namely *Acinetobacter* and *Paenibacilus*, which were not detected in the initial population, appeared to be positively affected by both pretreatments. It is being reported [43,44] that *Acinetobacter* strains are related to alcohol consumption as they can express alcohol dehydrogenase (ADH) to convert ethanol into acetate, reverting the solventogenic pathway 6. As shown in Figure 2, this reversed pathway probably not occurred during the essays since in all experiments, in their beginning and ending, ΔG_r_ for pathway 6 were thermodynamically feasible.

Conversely, ADH is also a molecule that is strongly related to bacterial quorum sensing with a key role in biofilm formation [44,45]. The growth of *Acinetobacter* was more stimulated in the thermal pretreatment, so this quorum sense mechanism could be linked with the best performance of this pretreatment to produce alcohols, considering that the presence of ethanol could have stimulated the expression of ADH, although the alcohol degradation metabolism was probably shut down. Acidic-thermal pretreatment preferentially stimulated the growth of *Paenibacilus*, which is directly related to alcohol production [46].

In addition to those changes, several other bacterial genera evolved differently depending on the pretreatment applied (Figure 9). The thermal pretreatment appeared to stimulate the growth of *Brevundimonas, Bacteroides, Butyrivibrio, Bacillus, Alcaligenes, Eubacterium, Clavibacter, Psychrobacter, Serratia* and *Microbulbifer*. The latter three being even exclusively detected in this pretreatment. On the opposite, the acidic-thermal pretreatment appeared to improve the growth of *Tissierella, Novosphingobium, Sedimentibacter* and *Yersinia*. The latter one being exclusively detected in this pretreatment.

The genus *Brevudimonas* started to increase in the thermal pretreatment after 8.3 days, coinciding with the very beginning of the exponential growth phase (Figure 4 and Table 5). Members belonging to this genus have already been shown to possess metabolic pathways related to the acidogenesis of alcohols [47]. High H_2_ partial pressures applied during the process might have rendered solventogenesis thermodynamically favourable through the reverse same metabolic pathway (Figure 3). The genus *Yersinia* started to increase in the acidic-thermal pretreatment after approximately 4.2 days, coinciding with the early exponential growth phase (Figure 4 and Table 5). Members belonging to this genus have already been shown to possess metabolic pathways to convert pyruvate into butanol and ethanol [48]. The genus *Sedimentibacter* is one of the predominant genus found in the fermentation of Baijiu, and it could be directly related to alcohol production [49]. It is noticeable that the genus *Clostridium*, which is composed of a high number of known alcohol producers [15,50–52], was detected for both pretreatments all along the process, without any significant variations. The genus *Clostridium* usually produces alcohol by converting glycerol and pyruvate into ethanol and butanol. Thus, this genus probably has an essential role in producing ethanol and butanol in both thermal and acidic-thermal pretreatments. There is no reporting on alcohol-producing metabolism for all other genera found within the experiments.

Other results were observed at higher taxonomic levels, notably decreasing microorganisms from the *Peptostreptococcaceae* family and the *Epsilonproteobacteria* class, in both pretreatments. *Peptostreptococcaceae* family represented almost half (45% of total bacterial OTUs) found in the initial inoculum (before any pretreatment), strongly decreased its presence to 5.4 ± 2.2% in thermal and 9.2 ± 3.1% in acidic-thermal pretreatments. *Epsilonproteobacteria* class represented up to 6% of total bacterial OTUs in the initial inoculum but significantly decreased to less than 0.3% of total bacterial OTUs in both thermal and acidic-thermal pretreatments. Such results indicate that neither bacteria from the *Peptostreptococcaceae* family nor *Epsilonproteobacteria* class played a significant role in the processes studied.

## 6 Conclusions

Characterization of the initial microbial population showed the presence of bacteria possessing metabolic pathways for alcohol production. Both thermal and acidic-thermal pretreatments were able to select the best adapted bacterial communities to produce ethanol and butanol. The significant decrease of members belonging to the family *Peptostreptococaceae* coupled to an increase of *Pseudomonas* appeared to be related to the best performance of ethanol and butanol production. The acidic-thermal pretreatment generated the best results, reaching a concentration of 87 mg L^-1^ of ethanol and of 143 mg L^-1^ of butanol after 240 hours of processing. The thermal pretreatment achieved the highest ethanol production (122 mg L^-1^) but at a much slower rate (after 710 hours). The two other pretreatments studied showed either instability and inconsistency for the thermal-acidic pretreatment or very low alcohol production for the acidic pretreatment. Among the VFAs added to the medium, butyrate was used solely as the substrate, acetate being produced along the process. The mass balance study highlighted another substrate input, which was probably originating from the inoculum. Thermodynamic data indicated that homoacetogenesis was a critical pathway to produce acetate from dissolved CO_2_ and H_2_. Finally, H_2_ partial pressure was a preponderant factor for solventogenesis, enabling alcohol production and inhibiting its consumption.

## 7 List of Abbreviations Used

STP: Standard temperature and pressure (0° C and 1 atm);
ppH_2_: Partial H_2_ pressure (atm);
R^2^: Levenberg-Marquardt algorithm correlation coefficient;
TVS: Total volatile solids (mg L^-1^);
OTU: Operational taxonomic unit;
HPLC: High performance liquid chromatography;
PCR: Polymerase chain reaction;
RDP: Ribosomal data project;
VFA: Volatile fatty acid;
But^-^: Butyrate;
Prop^-^: Propionate;
Ac^-^: Acetate;
But(OH): Butanol;
Prop(OH): Propanol;
Eth(OH): Ethanol;
*C*(*t*): Concentration in respect of time (mg L^-1^);
*t*: Time (h);
*t*_*i*_: Initial time for exponential growth phase (h);
*t*_*e*_: Ending time for exponential growth phase (h);
*t*_*m*_: Time in which maximum production rate (*r*_*max*_) is achieved (h);
*r*_*max*_: Maximum production rate (mg L^-1^ h^-1^);
*C*_*max*_: Maximum concentration (mg L^-1^);
ΔG_r_: Variation of Gibbs’ free energy at a given condition (kJ mol^-1^);
ΔG°_r_: Variation of Gibbs’ free energy at standard condition (kJ mol^-1^);
ΔG^F^_r_: Variation of Gibbs’ free energy at final experiment condition (kJ mol^-1^);
ΔG^I^_r_: Variation of Gibbs’ free energy at initial experiment condition (kJ mol^-1^).

## 9 Authors’ Contribution

All authors have planned the experiments. GM performed all the experiments, data treatment, statistics, and mathematical modeling. GB treated and analysed all sequencing data. All authors have analysed and discussed the results. GM have written the paper. All authors have read, reviewed, and approved the final manuscript. Experiments were conducted in Montréal branch of National Research Council of Canada from January to July of 2014.

## 10 Authors’ Informations

GM is professor of sanitation and applied microbiology of the Agricultural Engineering College at Campinas State University (FEAGRI/UNICAMP). This research was developed as an invited researcher at National Research Council of Canada (NRC) from October/2013 to October/2014, under a post-doctoral program in biofuels in the São Carlos Engineering School at University of São Paulo (EESC/USP).

## 11 Acknowledgements

This work was supported by FAPESP – Fundação de Amparo a Pesquisa do Estado de São Paulo (processes 2010/18.463-9 and 2013/18.172-2 – G. Mockaitis and 2009/15.984-0 – M. Zaiat), and the NRC – National Research Council of Canada (project A1-004645 – S.R. Guiot).

The authors thankfully acknowledge Marie-Josée Lévesque, Christine Maynard and Sylvie Sanschagrin for their assistance with the biomolecular techniques (DNA extraction, purification and PCR) and sequencing (through Ion Torrent™); and also, Stephane Deschamps and Alain Corriveau for their valuable contribution with physicochemical analysis (HPLC and gas chromatography of alcohols).

## 13 One-Sentence Summary

The inoculum pretreatment acid and thermal improved significantly the production of both ethanol and butanol, through an anaerobic solventogenic process, using acetate and butyrate as carbon source and H_2_ as electron donor.

## References

[1] Jones DT, Woods DR. Acetone-butanol fermentation revisited. Microbiol Rev 1986;50:484–524.

[2] Das D, Veziroglu T. Advances in biological hydrogen production processes. Int J Hydrogen Energy 2008;33:6046–57. https://doi.org/10.1016/j.ijhydene.2008.07.098.

[3] Baumann I, Westermann P. Microbial Production of Short Chain Fatty Acids from Lignocellulosic Biomass: Current Processes and Market. Biomed Res Int 2016;2016. https://doi.org/10.1155/2016/8469357.

[4] Levin D. Biohydrogen production: prospects and limitations to practical application. Int J Hydrogen Energy 2004;29:173–85. https://doi.org/10.1016/S0360-3199(03)00094-6.

[5] Steinbusch KJJ, Hamelers HVM, Buisman CJN. Alcohol production through volatile fatty acids reduction with hydrogen as electron donor by mixed cultures. Water Res 2008;42:4059–66. https://doi.org/10.1016/j.watres.2008.05.032.

[6] Zverlov V V, Berezina O, Velikodvorskaya G a, Schwarz WH. Bacterial acetone and butanol production by industrial fermentation in the Soviet Union: use of hydrolyzed agricultural waste for biorefinery. Appl Microbiol Biotechnol 2006;71:587–97. https://doi.org/10.1007/s00253-006-0445-z.

[7] Agler MT, Wrenn B a, Zinder SH, Angenent LT. Waste to bioproduct conversion with undefined mixed cultures: the carboxylate platform. Trends Biotechnol 2011;29:70–8. https://doi.org/10.1016/j.tibtech.2010.11.006.

[8] Dürre P. New insights and novel developments in clostridial acetone/butanol/isopropanol fermentation. Appl Microbiol Biotechnol 1998:639–48.

[9] Brynjarsdottir H, Wawiernia B, Orlygsson J. Ethanol Production from Sugars and Complex Biomass by Thermoanaerobacter AK 5□: The Effect of Electron-Scavenging Systems on End-Product Formation. Energy and Fuels 2012;26:4568–74.

[10] Jessen JE, Orlygsson J. Production of ethanol from sugars and lignocellulosic biomass by Thermoanaerobacter J1 isolated from a hot spring in Iceland. J Biomed Biotechnol 2012;2012:186982. https://doi.org/10.1155/2012/186982.

[11] Almarsdottir AR, Sigurbjornsdottir MA, Orlygsson J. Effect of various factors on ethanol yields from lignocellulosic biomass by Thermoanaerobacterium AK□□. Biotechnol Bioeng 2012;109:686–94. https://doi.org/10.1002/bit.24346.

[12] Crespo CF, Badshah M, Alvarez MT, Mattiasson B. Ethanol production by continuous fermentation of D-(+)-cellobiose, D-(+)-xylose and sugarcane bagasse hydrolysate using the thermoanaerobe Caloramator boliviensis. Bioresour Technol 2012;103:186–91. https://doi.org/10.1016/j.biortech.2011.10.020.

[13] Xu L, Tschirner U. Improved ethanol production from various carbohydrates through anaerobic thermophilic co-culture. Bioresour Technol 2011;102:10065–71. https://doi.org/10.1016/j.biortech.2011.08.067.

[14] Mes-Hartree M, Saddler J. Butanol production of Clostridium acetobutylicum grown on sugars found in hemicellulose hydrolysates. Biotechnol Lett 1982;4:247–52.

[15] Maddox I. Production of ethanol and n-butanol from hexose/pentose mixtures using consecutive fermentations with Saccharomyces cerevisiae and Clostridium acetobutylicum. Biotechnol Lett 1982;4:23–8.

[16] Ounine K, Petitdemange H, Raval G, Gay R. Acetone-butanol production from pentoses by Clostridium acetobutylicum. Biotechnol Lett 1983;5:605–10.

[17] Li Z, Shi Z, Li X. Models construction for acetone-butanol-ethanol fermentations with acetate/butyrate consecutively feeding by graph theory. Bioresour Technol 2014;159:320–6. https://doi.org/10.1016/j.biortech.2014.02.095.

[18] Kumar M, Goyal Y, Sarkar A, Gayen K. Comparative economic assessment of ABE fermentation based on cellulosic and non-cellulosic feedstocks. Appl Energy 2012;93:193–204. https://doi.org/10.1016/j.apenergy.2011.12.079.

[19] Kleerebezem R, van Loosdrecht MCM. Mixed culture biotechnology for bioenergy production. Curr Opin Biotechnol 2007;18:207–12. https://doi.org/10.1016/j.copbio.2007.05.001.

[20] Puig S, Coma M, Monclús H, van Loosdrecht MCM, Colprim J, Balaguer MD. Selection between alcohols and volatile fatty acids as external carbon sources for EBPR. Water Res 2008;42:557–66. https://doi.org/10.1016/j.watres.2007.07.050.

[21] O-Thong S, Prasertsan P, Birkeland N-K. Evaluation of methods for preparing hydrogen-producing seed inocula under thermophilic condition by process performance and microbial community analysis. Bioresour Technol 2009;100:909–18. https://doi.org/10.1016/j.biortech.2008.07.036.

[22] Luo G, Karakashev D, Xie L, Zhou Q, Angelidaki I. Long-term effect of inoculum pretreatment on fermentative hydrogen production by repeated batch cultivations: homoacetogenesis and methanogenesis as competitors to hydrogen production. Biotechnol Bioeng 2011;108:1816–27. https://doi.org/10.1002/bit.23122.

[23] Pendyala B, Chaganti SR, Lalman J a., Shanmugam SR, Heath DD, Lau PCK. Pretreating mixed anaerobic communities from different sources: Correlating the hydrogen yield with hydrogenase activity and microbial diversity. Int J Hydrogen Energy 2012;37:12175–86. https://doi.org/10.1016/j.ijhydene.2012.05.105.

[24] Kumar M, Gayen K, Saini S. Role of extracellular cues to trigger the metabolic phase shifting from acidogenesis to solventogenesis in Clostridium acetobutylicum. Bioresour Technol 2013;138:55–62. https://doi.org/10.1016/j.biortech.2013.03.159.

[25] Angelidaki I, Petersen SP, Ahring BK. Applied Microbiolog. v Effects of lipids on thermophilic anaerobic digestion and reduction of lipid inhibition upon addition of bentonite 1990:469–72.

[26] Sander R. Compilation of Henry ‘ s Law Constants for Inorganic and Organic Species of Potential Importance in Environmental Chemistry 1999.

[27] Lide DR, Frederikse HPR, editors. CRC Handbook of Chemistry and Physics. 76th ed. Boca Ratón, FL, USA: CRC Press Inc.; 1995.

[28] APHA. Standard methods for the examination of water and wastewater. 21st ed. Washington DC: 2005.

[29] Dilallo R, Albertson O. Volatile Acids by Direct Titration. J Water Pollut Control Fed 1961;33:356–65.

[30] Ripley L, Boyle W, Convrese J. Improved alkalimetric monitoring for anaerobic digestion of high-strength wastes. J Water Pollut Control Fed 1986;58:406–11.

[31] Guiot SR, Cimpoia R, Carayon G. Potential of wastewater-treating anaerobic granules for biomethanation of synthesis gas. Environ Sci Technol 2011;45:2006–12. https://doi.org/10.1021/es102728m.

[32] Muyzer G, Dewaal EC, Uitterlinden AG. Profiling of complex microbial populations by desnaturing gradient gel electrophoresis analysis ofpolymerase chain reaction amplified genes coding for 16S ribosomal RNA. Appl Environ Microbiol 1993;59:695–700.

[33] Griffiths RI, Whiteley a S, O’Donnell a G, Bailey MJ. Rapid method for coextraction of DNA and RNA from natural environments for analysis of ribosomal DNA- and rRNA-based microbial community composition. Appl Environ Microbiol 2000;66:5488–91.

[34] Lévesque MJ, La Boissière S, Thomas JC, Beaudet R, Villemur R. Rapid method for detecting Desulfitobacterium frappieri strain PCP-1 in soil by the polymerase chain reaction. Appl Microbiol Biotechnol 1997;47:719–25.

[35] Berthelet M, Whyte LG, Greer CW. Rapid, direct extraction of DNA from soils for PCR analysis using polyvinylpolypyrrolidone spin columns. FEMS Microbiol Lett 1996;138:17–22.

[36] Wang Q, Garrity GM, Tiedje JM, Cole JR. Naive Bayesian classifier for rapid assignment of rRNA sequences into the new bacterial taxonomy. Appl Environ Microbiol 2007;73:5261–7. https://doi.org/10.1128/AEM.00062-07.

[37] Claesson MJ, O’Sullivan O, Wang Q, Nikkilä J, Marchesi JR, Smidt H, et al. Comparative analysis of pyrosequencing and a phylogenetic microarray for exploring microbial community structures in the human distal intestine. PLoS One 2009;4:e6669. https://doi.org/10.1371/journal.pone.0006669.

[38] Mavrovouniotis ML. Group contributions for estimating standard gibbs energies of formation of biochemical compounds in aqueous solution. Biotechnol Bioeng 1990;36:1070–82. https://doi.org/10.1002/bit.260361013.

[39] Mavrovouniotis ML. Errata Group Contributions for Estimating Standard Gibbs Energies of Formation of Biochemical Compounds in Aqueous Solution 1991;38:803–4.

[40] Harper SR, Pohland FG. Recent Developments in Hydrogen Management During Anaerobic Biological Wastewater Treatment. Biotechnol Bioeng 1985;28:585–602.

[41] Speece RE. Anaerobic biotechnology for industrial wastewater. Vanderbilt University; 1996.

[42] Caspi R, Billington R, Fulcher CA, Keseler IM, Kothari A, Krummenacker M, et al. The MetaCyc database of metabolic pathways and enzymes. Nucleic Acids Res 2018;46:D633–9. https://doi.org/10.1093/nar/gkx935.

[43] John J, Saranathan R, Adigopula LN, Thamodharan V, Singh SP, Pragna Lakshmi T, et al. The quorum sensing molecule N-acyl homoserine lactone produced by Acinetobacter baumannii displays antibacterial and anticancer properties. Biofouling 2016;32:1029–47. https://doi.org/10.1080/08927014.2016.1221946.

[44] Zhang K, Yang X, Yang J, Qiao X, Li F, Liu X, et al. Alcohol dehydrogenase modulates quorum sensing in biofilm formations of Acinetobacter baumannii. Microb Pathog 2020;148:104451. https://doi.org/10.1016/j.micpath.2020.104451.

[45] Subhadra B, Hwan Oh M, Hee Choi C. Quorum sensing in *Acinetobacter*: with special emphasis on antibiotic resistance, biofilm formation and quorum quenching. AIMS Microbiol 2016;2:27–41. https://doi.org/10.3934/microbiol.2016.1.27.

[46] Grady EN, MacDonald J, Liu L, Richman A, Yuan ZC. Current knowledge and perspectives of Paenibacillus: A review. Microb Cell Fact 2016;15:203. https://doi.org/10.1186/s12934-016-0603-7.

[47] Leifson E, Hugh R. A new type of polar monotrichous flagellation. J Gen Microbiol 1954;10:68–70. https://doi.org/10.1099/00221287-10-1-68.

[48] Caspi R, Altman T, Dale JM, Dreher K, Fulcher CA, Gilham F, et al. The MetaCyc database of metabolic pathways and enzymes and the BioCyc collection of pathway/genome databases. Nucleic Acids Res 2010;38:D473–9. https://doi.org/10.1093/nar/gkp875.

[49] Xu J, Sun L, Xing X, Sun Z, Gu H, Lu X, et al. Culturing Bacteria From Fermentation Pit Muds of Baijiu With Culturomics and Amplicon-Based Metagenomic Approaches. Front Microbiol 2020;11:1223. https://doi.org/10.3389/fmicb.2020.01223.

[50] Tracy BP, Jones SW, Fast AG, Indurthi DC, Papoutsakis ET. Clostridia: the importance of their exceptional substrate and metabolite diversity for biofuel and biorefinery applications. Curr Opin Biotechnol 2012;23:364–81. https://doi.org/10.1016/j.copbio.2011.10.008.

[51] Ueki A, Hirono T, Sato E, Mitani A, Ueki K. Ethanol and amylase production by a newly isolated Clostridium sp. World J Microbiol … 1991;7:385–93.

[52] Bruant G, Lévesque M-J, Peter C, Guiot SR, Masson L. Genomic analysis of carbon monoxide utilization and butanol production by Clostridium carboxidivorans strain P7. PLoS One 2010;5:e13033. https://doi.org/10.1371/journal.pone.0013033.

