## Supplementary Information for "Microbial Communities Performing Hydrogen Solventogenic Metabolism of Volatile Fatty Acids"

### 1.1 Modified Boltzmann Sigmoidal Equation

This addendum has the objective to explain all the steps to achieve the modified Boltzmann sigmoidal equation as presented in Equation 1. In addition to this, all the assumption to infer the starting and ending points of the exponential growth phase, shown by Equations 2 and 3 will be explained as well.

The sigmoidal is the standard shape of a bacterial monophasic growth curve. For applying such model, it was assumed that microbial growth could be linearly associated with both substrates consumption and products production time profiles. Thus, the sigmoidal function was chosen to describe the time profile of ethanol and butanol production, and it was modified so its parameters could mean a biological significance.

The original form of sigmoidal equation proposed by Boltzmann in 1879 is a logistic function modification, and is shown by Equation A.

| $y(x)=y_{i}-\frac{(y_{i}-y_{f})}{1+e^{\left( \frac{x-x_{0}}{\gamma} \right)}}$ | Equation A |
| --- | --- |

Where $y_{f}$ is the maximum value reached by the ordinate, defined by $y_{f}=\lim_{x\to+\infty} y(x)$; $y_{i}$ is the minimum value reached by the ordinate, defined by $y_{i}=\lim_{x\to-\infty} y(x)$; $x_{0}$ is the abscissa value when the slope value is maximum, defined by $x_{0}=x \Longleftrightarrow y\left( x_{0} \right)=\frac{y_{f}-y_{i}}{2}$; and $\gamma$ is an slope correlated value which describes its behavior.

For applying this mathematical model for describing more adequately the behavior of a microbiological system, it was considered some boundary conditions. Since this model was applied for alcohol production evaluation, and none of these products was detected at the beginning of the experiments (when time = 0), the parameter $y_{i}$ was considered null (i.e. $y_{i}=0$). Also, in spite of the value of the $\gamma$ parameter could be related with the slope of the sigmoidal function, it does not represent any kind of comparable parameter. In this sense, this parameter was replaced by the value of the maximum slope of the sigmoidal function, determined as shown by Equation B.

| $r(x)=\frac{dy(x)}{dx}=\frac{y_{f}}{\gamma}\cdot\frac{e^{\left( \frac{x-x_{0}}{\gamma} \right)}}{\left[ 1+e^{\left( \frac{x-x_{0}}{\gamma} \right)} \right]^{2}}$ | Equation B |
| --- | --- |

The maximum value of the function slope is reached when $x=x_{0}$, which simplifies the Equation B into the Equation C.

| $r_{max}=r(x_{0})=\frac{y_{f}}{4\cdot\gamma}$ | Equation C |
| --- | --- |

Replacing the Equation C in the Equation A, also constraining the value $y_{i}=0$ as boundary condition, it is possible to rewrite the Equation A in terms of $r_{max}$, as depicted in Equation D, which is the modified Boltzmann equation described as Equation 1.

| $y(x)=y_{f}-\frac{y_{f}}{1+e^{\left( \frac{4{\cdot r}_{max}(x-x_{0})}{y_{f}} \right)}}$ | Equation D |
| --- | --- |

The Figure A shows the Equation D and its derivative for arbitrary parameters values, and places the parameters in the graph.

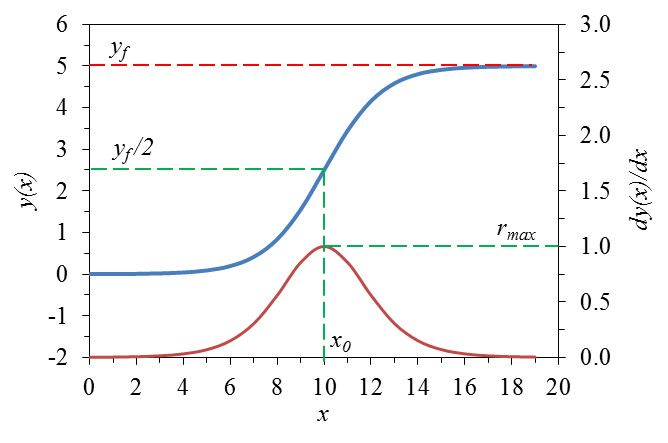

Figure A – Shape of the modified Boltzmann equation (blue filled line, $y(x)$) and its derivative (red filled line, $dy(x)/dx$)), for $y_{f}=5$; $x_{0}=10$ and $r_{max}=1$. The dashed red line shows the value of $y_{f}$, as the dashed green line shows $x_{0}$ in $y(x)$ and $r_{max}$ for $dy(x)/dx$.

Throughout this modified equation, it was possible to determine, only by evaluating its parameters, the maximum production of ethanol and butanol ($y_{f}$), its maximum rate ($r_{max}$) and the time in which this maximum rate was achieved ($x_{0}$).

The initial and final points of the exponential phase ($x_{i}$ and $x_{f}$) are also important parameters to compare and evaluate the process. These values were estimated approximating the shape of the exponential phase to a straight-line equation, in which the parameter $r_{max}$ was the angular coefficient. Thus, the behavior of $y(x)$ in between the interval [$x_{i},x_{f}$] was modeled as a function $y^{*}(x)$, described by the Equation E, only for means to calculate both $x_{i}$ and $x_{f}$.

| $y^{*}\left( x \right)=r_{max}\cdot\left( x-x_{0} \right)+\frac{y_{f}}{2}$ | Equation E |
| --- | --- |

Thus, the parameters $x_{i}$ and $x_{f}$ are calculated equating the Equation E to 0 and $y_{f}$, respectively, leading then to the expressions represented for the Equations F and G.

| $y^{*}\left( x_{i} \right)=0\Rightarrow x_{i}=x_{0}-\frac{y_{f}}{2\cdot r_{max}}$ | Equation F |
| --- | --- |
| $y^{*}\left( x_{f} \right)=y_{f}\Rightarrow x_{f}=x_{0}+\frac{y_{f}}{2\cdot r_{max}}$ | Equation G |

These parameters shall be considered as an estimation of the initial and final time of the exponential growth phase. A graphic explanation of this method is depicted on Figure B.

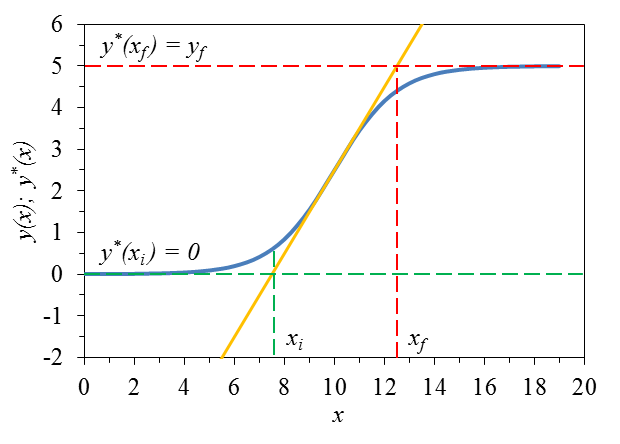

Figure B – Estimation of initial ($x_{i}$) and final ($x_{f}$) time of the exponential growth phase. The Equation E function (yellow filled line, $y^{*}(x))$ and the modified Boltzmann equation (blue filled line, $y(x)$) are shown. The green and the red dashed lines show the boundary condition for calculation of $x_{i}$ and $x_{f}$, respectively.

Finally, the time length of the exponential time could be obtained subtracting $x_{f}$ from $x_{i}$, leading to the Equation H.

| $x_{e}=x_{f}-x_{i}\Rightarrow x_{e}=\frac{y_{f}}{r_{max}}$ | Equation H |
| --- | --- |
